# Combined prophylactic and therapeutic use maximizes hydroxychloroquine anti-SARS-CoV-2 effects *in vitro*

**DOI:** 10.1101/2020.03.29.014407

**Authors:** Nicola Clementi, Elena Criscuolo, Roberta Antonia Diotti, Roberto Ferrarese, Matteo Castelli, Roberto Burioni, Massimo Clementi, Nicasio Mancini

## Abstract

While the SARS-CoV-2 pandemic is hardly hitting the world, it is of extreme importance that significant *in vitro* observations guide the quick set up of clinical trials. In this study, we evidence that the anti-SARS-CoV2 activity of a clinically achievable hydroxychloroquine concentration is maximized only when administered before and after the infection of Vero E6 cells. This strongly suggests that only a combined prophylactic and therapeutic use of hydroxychloroquine may be effective in limiting viral replication in patients.

## Introduction

On 11^th^ March 2020, WHO Director-General characterized COVID-19 as a pandemic. As of 27^th^ March 2020, 549,604 total cases were confirmed, accounting for 24,863 deaths ^1^. To date, the clinical management of COVID-19 subjects almost exclusively consists of supportive therapy and, in particular, of ventilatory support for the most severe cases. Several drugs, some already in clinical trials, are currently being used in order to limit the inflammatory response or to hamper SARS-CoV-2 replication. Regarding the latter, no SARS-CoV-2-specific drugs are currently available and, as a consequence, there is the need for repurposing drugs used in other settings, such as chloroquine (CQ) and hydroxychloroquine (HCQ) ^2,3^. Several studies already demonstrated the *in vitro* efficacy of CQ and HCQ, evidencing the latter’s higher activity on SARS-CoV-2 ^4,5^. Importantly, a recent study also modeled the main pharmacokinetic features of HCQ trying to infer its lung concentration after different dosing regimens^6^.

It is therefore important to use this data in order to set up possible scenarios related to the real clinical use of this drug. In this study, we tested HCQ against a SARS-CoV-2 Italian clinical isolate, by using different protocols of drug administration corresponding to its possible prophylactic, therapeutic and prophylactic/therapeutic use in patients. A single HCQ concentration easily reachable in the lung and characterized by high anti-SARS-CoV-2 activity was used.

## Methods

### Cells, virus and antivirals

Vero E6 (Vero C1008, clone E6 – CRL-1586; ATCC) cells were cultured in Dulbecco’s Modified Eagle Medium (DMEM) supplemented with non-essential amino acids (NEAA), penicillin/streptomycin (P/S), Hepes buffer and 10% (v/v) Fetal bovine serum (FBS). A clinical isolate hCoV-19/Italy/UniSR1/2020 (GISAID accession ID: EPI_ISL_413489) was isolated and propagated in Vero E6 cells, and viral titer was determined by 50% tissue culture infective dose (TCID_50_) and plaque assay for confirming the obtained titer. All the infection experiments were performed in a biosafety level-3 (BLS-3) laboratory of Microbiology and Virology at Vita-Salute San Raffaele University, Milan, Italy. Bafilomycin A1 (BFLA) was obtained from Merck, Hydroxychloroquine (HCQ) was obtained from Sigma-Aldrich.

### Virus isolation

An aliquot (0.8 mL) of the transport medium of the nasopharyngeal swab (COPAN’s kit UTM® universal viral transport medium - COPAN) of a mildly symptomatic SARS-CoV-2 infected patient was mixed with an equal volume of DMEM without FBS and supplemented with double concentration of P/S and Amphotericin B. The mixture was added to 80% confluent Vero E6 cells monolayer seeded into a 25 cm^2^ tissue culture flask. After 1 h adsorption at 37°C, 3 mL of DMEM supplemented with 2% FBS and Amphotericin B were added. Twenty-four hours post-infection (hpi) another 2 mL of DMEM supplemented with 2% FBS and Amphotericin B were added. Live images were acquired (Olympus CKX41 inverted phase-contrast microscopy) daily for evidence of cytopathic effects (CPE), and aliquots were collected for viral RNA extraction and In-house one-step real-time RT-PCR assay (10.1016/ S0140-6736(20)30154-9). Five days post-infection (dpi) cells and supernatant were collected, aliquoted and stored at −80°C (P1). For secondary (P2) virus stock, Vero E6 cells seeded into 25 cm^2^ tissue culture flasks were infected with 0.5 mL of P1 stored aliquot, and infected cells and supernatant were collected 48 hpi and stored at −80°C. For tertiary (P3) virus stock, Vero E6 cells seeded into 75 cm^2^ tissue culture flasks were infected with 1.5 mL of P2 stored aliquot and prepared as above described.

### Virus titration

P3 virus stocks were titrated using both Plaque Reduction Assay (PRA, PFU/mL) and Endpoint Dilutions Assay (EDA, TCID_50_/mL). For PRA, confluent monolayers of Vero E6 cells were infected with 10-fold-dilutions of virus stock. After 1 h of adsorption at 37°C, the cell-free virus was removed. Cells were then incubated for 46 h in DMEM containing 2% FBS and 0.5% agarose. Cells were fixed and stained, and viral plaques were counted. For EDA, Vero E6 cells (4 × 10^5^ cells/mL) were seeded into 96 wells plates and infected with base 10 dilutions of virus stock. After 1 h of adsorption at 37°C, the cell-free virus was removed, and complete medium was added to cells. After 48 h, cells were observed to evaluate CPE. TCID_50_/mL was calculated according to the Reed–Muench method.

### Sequence analysis

Viral genome from supernatant infected cells was extracted using QIAamp Viral RNA Mini Kit following manufacturers’ instructions. Reverse transcription and subsequent amplification were performed using random hexamer primers. The amplicons were sequenced on Illumina MiSeq NGS platform (Illumina, San Diego, CA, USA). Amplicon purification and quantification were performed by Agencourt AMPure XP (Beckman Coulter, Villepinte, France) and Qubit dsDNA Assay Kit (ThermoFisher Scientific, Waltham, MA, USA) respectively. Library preparation was performed by using Nextera XT DNA Library Prep Kit (Illumina, San Diego, CA, USA). The library generated was then diluted and sequenced with MiSeq Reagent Kit v2 (300-cycles) (Illumina, San Diego, CA, USA) on MiSeq platform. The quality of raw sequences obtained from MiSeq run was first checked using FastQC (v 0.11.5) (Babraham Bioinformatics). The reads were aligned on reference sequence (GISAID accession ID: EPI_ISL_412973) using BWA-mem and rescued using Samtools alignment/Map (v 1.9) and bamtoFastq. Finally, the contigs were generated using SPAdes (v 3.12.0).

### Time-of-addition experiments of HCQ

Vero E6 cells (4 × 10^5^ cells/mL) were seeded into 96 wells plates and treated with HCQ (10 μM) at different stages of virus infection. For *full-time* treatment, cells were pre-treated with the drug for 1 h prior to virus infection at 37°C, followed by virus adsorption for 1 h in the presence of the molecule. Then, cells were washed with PBS, and further cultured at 37°C with the molecule-containing medium until the end of the experiment. For *pre-adsorption* treatment, the agent was added to the cells for 1 h at 37°C before virus infection and maintained during virus adsorption. Then, the mixture was replaced with fresh medium without molecule till the end of the experiment. For *post-adsorption* assays, the drug-containing medium was added to cells only after virus adsorption and maintained until the end of the experiment. Then, cells were washed with PBS, and further cultured at 37°C with the molecule-containing medium until the end of the experiment. BFLA (100 nM) was tested as control of inhibition of viral infectivity at a phagolysosome level only in a *pre-adsorption* treatment, alone or in combination with *pre-adsorption* HCQ treatment. Uninfected cells were included in all experimental settings to exclude possible drug-toxicity CPE. For all the experimental groups, cells were infected with 50 TCID_50_/mL (98 PFU/mL) SARS-CoV-2 and absorption was performed for 1h at 37°C or 4°C. Live images were acquired (Olympus CKX41 inverted phase-contrast microscopy) and cell supernatants were collected for real-time PCR (RT PCR) analysis at 72 hpi. All conditions were tested in quadruplicate.

### Viral RNA extraction and real-time RT-PCR

Viral RNA was purified from 140 mL of cell culture supernatant using the QIAamp Viral RNA Mini Kit (QIAGEN), following the manufacturer’s instructions. Subsequently, the purified RNA was used to perform the synthesis of first-strand cDNA, using the SuperScript™ First-Strand Synthesis System for RT-PCR (Thermo Fisher Scientific), following the manufacturer’s instruction. Real-time PCR, using SYBR® Green dye-based PCR amplification and detection method, was performed in order to detect the cDNA. We used the SYBR™ Green PCR Master Mix (Thermo Fisher Scientific) the forward primer N2F: TTA CAA ACA TTG GCC GCA AA, the reverse primer N2R: GCG CGA CAT TCC GAA GAA, and the following PCR conditions: 95°C for 2 min, 45 cycles of 95°C for 20 sec, annealing at 55°C for 20 sec and elongation at 72°C for 30 sec, followed by a final elongation at 72°C for 10 min ^7^. RT-PCR was performed using the ABI-PRISM 7900HT Fast Real-Time instruments (Applied Biosystems) by using optical-grade 96-well plates. Samples were run in duplicate in a total volume of 20 μL.

### Statistical analysis

CPE observed for different experimental settings using HCQ and BFLA, alone or in combination, were normalized to corresponding virus infection control. RT-PCR results were analyzed calculating Delta (Δ) Ct as the difference between Ct values obtained for experimental settings and infection control. Then, two-way ANOVA and Tukey’s multiple comparisons test were performed for the evaluation of Ct differences (GraphPad Prism 8).

## Results

### Virus isolation and sequencing

Virus isolation was achieved after less than 72 hours. At 48 hpi the cytopathic effect (CPE) was already evident on Vero E6 cells. NGS analysis was performed by Illumina MiSeq obtaining the whole genome sequence of the cultured isolate hCoV-19/Italy/UniSR1/2020 (GISAID accession ID: EPI_ISL_413489).

### Time-of-addition experiments of HCQ and BFLA

The two molecules were tested using different experimental protocols, and virus adsorption was performed at both 37°C and 4°C. CPE was assessed at 72 hpi (**Fig. 1 and 2**). Virus infection positive control showed marked effects on cell morphology at 37°C as well as 4°C adsorption conditions. HCQ was effective in *full-time* treatment at both adsorption temperatures, and in *post-adsorption* treatment only when the virus was added to cells for 1 h at 4°C. The molecules did not show the same degree of protection from CPE in *pre-adsorption* treatment at both adsorption temperatures, as well as in *post-adsorption* treatment at 37°C adsorption. BFLA was tested as control of inhibition of viral infectivity at a phagolysosome level in a *pre-adsorption* treatment, showing full CPE protection at 37°C virus adsorption, while only partial protection was observed when the virus was added to cells at 4°C (**Fig. 2**). No drug-related cytotoxic effect was observed on uninfected cells, in all experimental settings (data not shown).

**Fig. 1.**
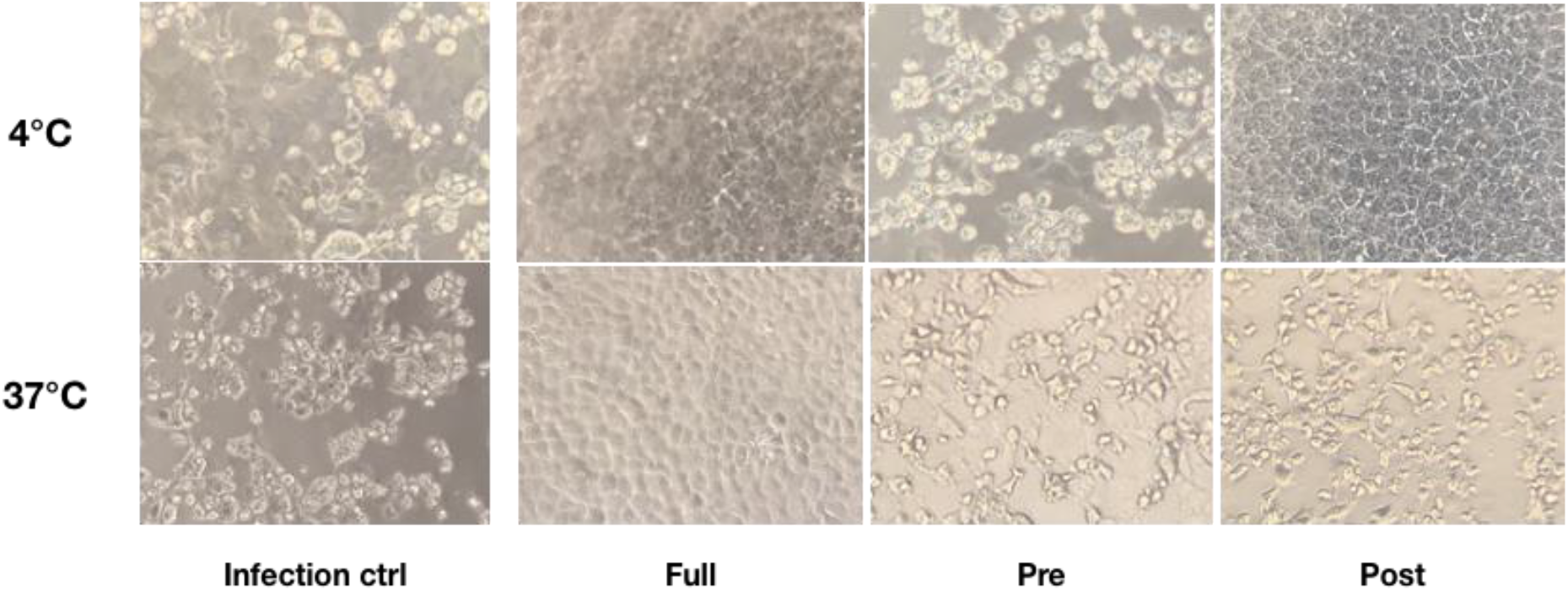
CPE on infected cells treated with HCQ in different experimental settings. Bright-field microscopy images (20x magnification, 72 hpi) of representative CPE of hCoV-19/Italy/UniSR1/2020 (GISAID accession ID: EPI_ISL_413489) isolate detected on both untreated cells (Infection control) and treated cells with different experimental settings of HCQ treatment. Virus adsorption was performed at 4°C and 37°C.

**Fig. 2.**
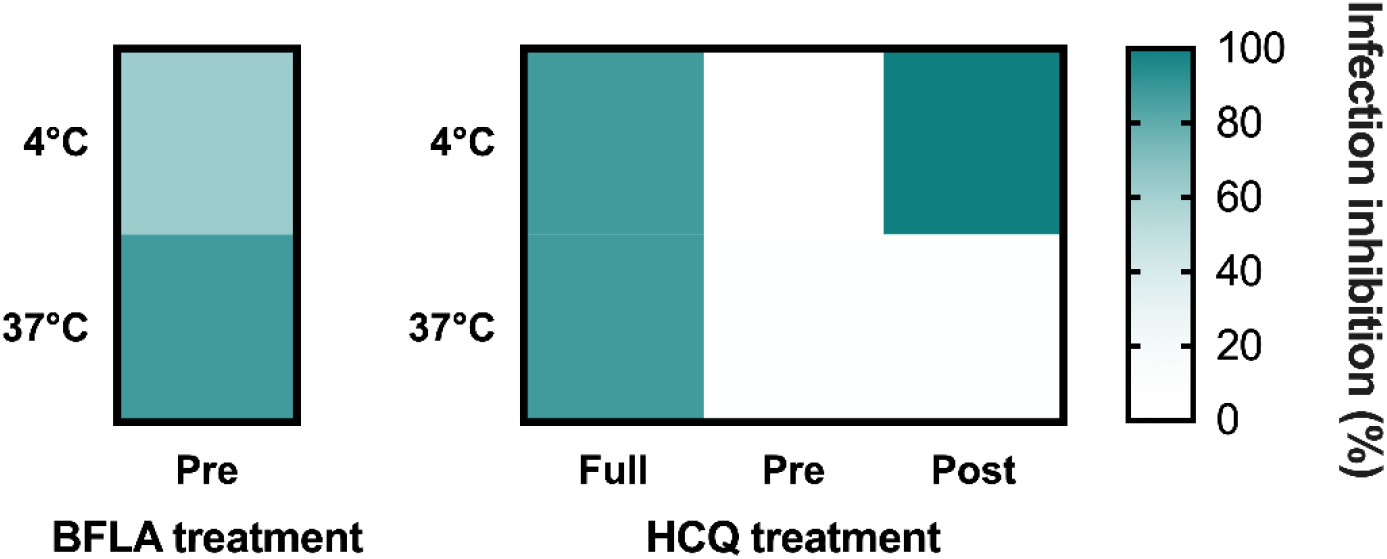
HCQ CPE reduction on VeroE6 cells infected with SARS-CoV-2. Green gradient indicates the reciprocal of CPE detected in treated cells compared to virus infection control (white color corresponds to 100% CPE). Protection levels are indicated by color gradient: white corresponds to no protection, dark blue shows full protection. BFLA treatments are reported as experimental control of virus fusion process inhibition. Virus adsorption was performed at 4°C and 37°C.

### RT-PCR analysis

Cell supernatants of different experimental settings were collected and analyzed by RT-PCR, and results confirmed CPE data analysis (**Fig. 3**). In detail, a significant statistical difference of Delta Ct was observed with HCQ *full-time* treatment compared to infection control, both at 37°C (*P* < 0.05) and 4°C (*P* < 0.0001) virus adsorption, while HCQ *post-adsorption* treatment was effective (*P* < 0.0001) only when the virus was added to cells at 4°C. Interestingly, BFLA addition to HCQ *pre-adsorption* treatment resulted in a significant statistical difference of Delta Ct when compared to infection control at 37°C virus adsorption.

**Fig. 3.**
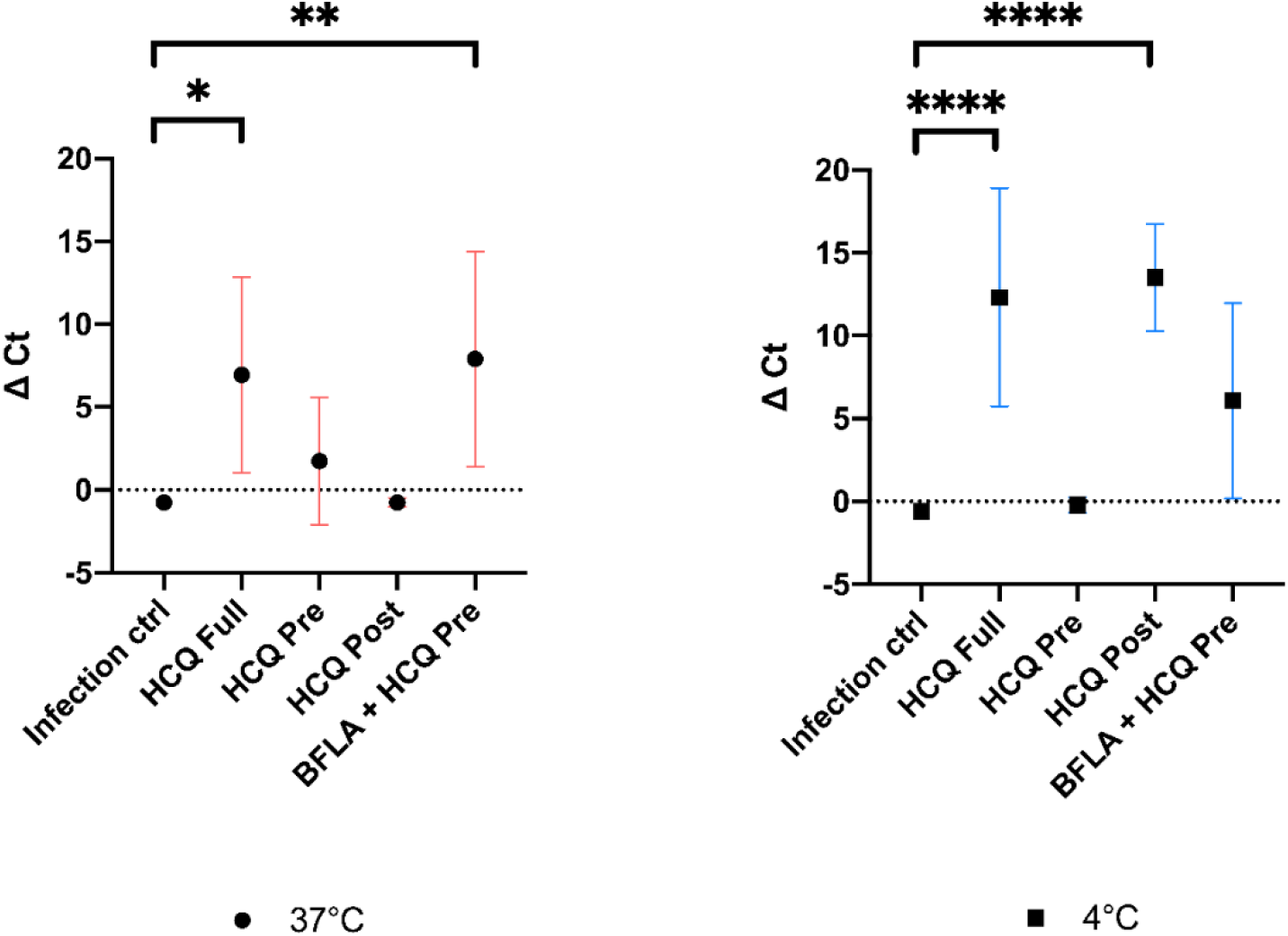
RT-PCR analysis of cell supernatants of different experimental settings. Ct levels are inversely proportional to the amount of target nucleic acid in the sample (the lower the Ct level the greater the amount of virus within the tested supernatant). Graphs show virus adsorption at 37°C and at 4°C. Delta (Δ) Ct are represented in y axis. Median values for all experimental replicates, tested each one in duplicate in RT-PCR, and 95% IC range reported with error bars (* *P* < 0.05; ** *P* < 0.01; **** *P* < 0.0001).

## Discussion

In the lack of SARS-CoV-2-specific drugs it is extremely important to evaluate the clinical potential of drug-repurposing in order to face the current pandemic ^8^. HCQ has gained the attention of the scientific and medical community based on previous in vitro data on similar viruses (SARS and MERS) and on preliminary reports discussing its possible clinical effectiveness ^2–4^. In this atypical context, in which on-field medicine often anticipates experimental laboratory pre-clinic, there is urgent need of prompt experiments addressing specific clinical questions. For example, it is not clear what is the best administration regimen to maximise possible HCQ anti-viral effectiveness in COVID-19 patients. At this regard, a recent study evaluated the direct anti-SARS-CoV-2 effect *in vitro* and, importantly, modelled its bioavailability at the lung level where it could maximally exert its antiviral activity. Based on a dosing regimen of 400 mg given twice daily for 1 day, followed by 200 mg twice daily for 4 more days, it also suggested the more useful drug concentrations to be used in clinically-oriented phenotypic laboratory assays^5,6,9^.

On that basis, we evaluated the HCQ antiviral activity when administered before (*pre-adsorption*), after (*post-adsorption*) or before and after (*full-time*) virus adsorption in order to simulate its possible prophylactic, therapeutic and prophylactic/therapeutic clinical use. Moreover, we focused our attention on a single concentration (10 uM) easily achieved, well tolerated and endowed with a strong antiviral activity^6^. Moreover, in order to speculate on its possible mechanism of action, we also evaluated HCQ activity performing virus adsorption at 37 and 4°C. In fact, at 37°C the virus enters the cell in a more physiological context while, conversely, at 4°C virus it can dock to the cell receptor, but its internalisation is much more limited.

In the prophylactic setting (*pre-adsorption*), 10uM HCQ did not interfere effectively with the viral replicative cycle neither at 37°C nor at 4°C, as evidenced in the CPE and the RT-PCR analysis. A limited antiviral activity was also observed in the therapeutic setting (*post-adsorption*) but, interestingly, a higher HCQ antiviral activity was observed at 4°C, suggesting its predominant SARS-CoV-2 interference at the endosomal level ^10,11^. Overall, and most importantly, these results suggest a limited activity of HCQ when administered only prophylactically or therapeutically.

On the contrary, a significant antiviral activity was observed in the prophylactic/therapeutic (*full-time*) experimental setting both at 37°C and 4°C, as evidenced both by CPE and RT-PCR analyses. This observation allows to speculate on the need of a combined prophylactic and therapeutic clinical use of HCQ on in order to maximise its antiviral effects. In the atypical scenario of an ongoing pandemic, pre-clinical medical research should be focused on simple and fast observations potentially useful for the prompt set up of clinical trials. As an example, our observation could be easily translated in a clinical study on extremely high-risk categories, such as health care workers, based on the prophylactic administration of HCQ followed by its therapeutic use in case of positivity to SARS-CoV-2. The time is crucial during a pandemic and the correct clinical repurposing of HCQ could be really invaluable.

